# Free-living amoebae act as transient permissive hosts for *Leptospira spp*

**DOI:** 10.64898/2026.03.16.712017

**Authors:** Adriana Luga, Catherine Inizan, Eva Meunier, Amélie Albon, Valérie Burtet-Sarramegna, Mathieu Picardeau, Cyrille Goarant, Roman Thibeaux

## Abstract

**Background:** Leptospirosis is a zoonotic disease caused by pathogenic *Leptospira* spp., which persist in soil and water environments for extended periods of time. The mechanisms enabling this environmental survival remain elusive. Free-living amoebae (FLA) are widespread protozoa that act as reservoirs or “Trojan horses” for numerous bacterial pathogens, protecting them from stress and contributing to their persistence. Whether pathogenic *Leptospira* exploit similar interactions with FLA has not been resolved.

**Methodology/Principal Findings:** Using live confocal microscopy, flow cytometry, and gentamicin protection assays, we investigated the interactions between pathogenic (*Leptospira interrogans*) and saprophytic (*Leptospira biflexa*) leptospires with three FLA species: *Acanthamoeba castellanii*, *Dictyostelium discoideum*, and *Hartmannella vermiformis*. While rapid internalization was observed, entry was only partially dependent on actin-driven processes and was enhanced by the presence of live bacteria. Following internalization, bacteria persisted for at least 48h as indicated by colony-forming assays. However, no evidence of intracellular replication was detected. The number of fluorescently labeled leptospires progressively declined over time, providing further evidence of leptospires survival without multiplication. Finally, analysis of environmental soils in New Caledonia showed co-occurrence of FLA and *Leptospira*. Soil-derived FLA also internalized pathogenic *Leptospira in vitro*, showing that these interactions extend to natural isolates.

**Conclusions/Significance:** Our results demonstrate that free-living amoebae internalize both pathogenic and saprophytic leptospires and allow their transient persistence without replication. By providing protection and prolonging viability in soil environments, FLA may contribute to the ecological maintenance of *Leptospira*. These findings pinpoint FLA as potential environmental reservoirs that could play a role in shaping leptospires survival strategies relevant for transmission and host infection.

**Author Summary:** For bacteria living in soils and freshwater environments, survival depends on their ability to adapt to complex ecological landscapes populated by numerous predators and competitors. In such habitats, interactions with other microorganisms are unavoidable and may shape long-term survival strategies. Pathogenic *Leptospira*, the bacteria responsible for leptospirosis, can persist for long periods outside their hosts, yet the ecological mechanisms supporting this environmental survival remain poorly understood. In soil and freshwater ecosystems, microscopic predators known as free-living amoebae commonly feed on bacteria. However, several bacterial pathogens can survive inside these amoebae and use them as temporary shelters. Because ancestral *Leptospira* were soil-dwelling saprophytes, interactions with amoebae likely represent an ancient ecological relationship in which successful survival strategies may have evolved and remain conserved in present-day pathogenic species.

With this perspective in mind, we used microscopy approaches and bacterial viability assays to investigate whether *Leptospira* interacts with amoebae. We found that several amoeba species rapidly engulf both pathogenic and non-pathogenic *Leptospira*. Once internalized, the bacteria remained viable for up to two days but did not multiply. We also detected both amoebae and *Leptospira* in the same soil samples and showed that environmental amoebae could internalize the bacteria. These findings suggest that amoebae may act as temporary shelters for *Leptospira*, helping them persist in soils and water and potentially contributing to the environmental stage of leptospirosis transmission.

## 1. Introduction

Leptospirosis is a globally distributed zoonosis caused by pathogenic *Leptospira* spp. These bacteria are transmitted to humans primarily through exposure to environments contaminated by the urine of infected reservoir hosts. Beyond their mammalian hosts, leptospires have also been detected and isolated from soils and water, supporting their classification as environmental bacteria. Under specific conditions such as water-saturated soils, experimental evidence indicates that leptospires, including pathogenic species, are able to not only persist but also multiply in the environment (1). Although considered fragile under laboratory conditions, early and recent studies have demonstrated their ability to persist for extended periods in natural settings (2), with a survival up to six months in water-saturated soils (3) and have been repeatedly detected at intervals of up to nine weeks in river soils (4). In addition, low bacterial load in soils has been proposed as sufficient to sustain human infection risk (5). Field and laboratory studies have therefore documented the repeated detection of pathogenic leptospires in these environments (4,6–10), consistent with an ecologically relevant extra-host phase. Mechanistic explanations for persistence include biofilm formation (11,12) and tolerance to physicochemical stress (13). However, the specific biotic interactions that enable survival, multiplication or concentration of leptospires in natural environments remain poorly explored (14).

Free-living amoebae (FLA) are ubiquitous protozoa in soil and freshwater systems recognized as important hosts for a broad range of human bacterial pathogens. *Acanthamoeba castellanii*, for example, can harbor *Helicobacter pylori*, which remains viable and resistant to chlorination following intracellular residence (15). Similarly, *Dictyostelium discoideum*, a professional phagocyte widely used as a biomedical model, is permissive to infection by *Legionella pneumophila*, *Mycobacterium tuberculosis*, and *Salmonella enterica* (16). *Hartmannella vermiformi*s is another environmentally widespread FLA consistently found in association with *L. pneumophila* and within which these bacteria replicate (17). These findings highlight the role of FLA as “Trojan horses” that may protect some pathogenic bacteria against environmental stresses thus enhancing their persistence and dispersal. Importantly, many FLA can differentiate into highly resistant cyst forms under adverse environmental conditions (18,19). Encystment provides protection against desiccation, nutrient deprivation, temperature fluctuations, and chemical stress, potentially offering intracellular bacteria an additional layer of environmental shielding.

Given their shared ecological niche, *Leptospira* and FLA are likely to interact in natural habitats. Yet, whether pathogenic *Leptospira* interact with FLA similarly to other human pathogens has not been assessed. This question is particularly relevant in light of recent evidence showing that pathogenic and saprophytic *Leptospira* can be internalized by mammalian macrophages without replicating (20). In these mammalian cells, bacterial load decreases over time, but both *L. interrogans* and *L. biflexa* can exit, suggesting survival strategies that allow transient persistence inside professional phagocytes. Although FLA are not immune cells and lack the receptor repertoire and immunological circuitry of mammalian macrophages, they share several conserved cellular modules relevant to phagocytosis, phagolysosomal processing, and antimicrobial responses (21,22). Comparative work has highlighted functional parallels between predatory amoebae and pro-inflammatory (M1-like) macrophage programs, supporting the view that environmental phagocytes can impose selective pressures that resemble, without recapitulating, those encountered in mammalian phagocytes (23). In this framework, free-living amoebae may represent ecological “training grounds” (24), in which traits promoting persistence within phagocytic cells could be selected and later prove advantageous during interactions with macrophages.

In this study, we investigated the interactions between pathogenic and saprophytic *Leptospira* and different FLA species, focusing on their trophozoite stage, the metabolically active and phagocytic form of amoebae, with the working hypothesis that FLA may act as transient or stable reservoirs. Specifically, we addressed three questions: (i) do leptospires associate with and become internalized by trophozoites? (ii) if leptospires are internalized, can they persist within amoebae as viable bacteria? and (iii) can these interactions be observed in environmental isolates of FLA and leptospires naturally co-occurring in soil? To answer these questions, we combined gentamicin protection assays with live confocal microscopy, flow cytometry, and - culturability-based readouts.

Our results show that both pathogenic *L. interrogans* and saprophytic *L. biflexa* are rapidly internalized by multiple FLA species, including *A. castellanii*, *D. discoideum*, and *H. vermiformis*. Internalization was only partially dependent on classical phagocytic pathways and was enhanced by bacterial viability, suggesting active bacterial contribution. Although intracellular leptospires remained viable for up to 48 h following internalization, no evidence of replication was observed. Finally, we isolated both *Leptospira* and FLA from soil samples in New Caledonia and demonstrated that soil-derived FLA also internalize leptospires, extending our *in vitro* observations with model organisms to a natural context. These findings support a model in which FLA could transiently contribute to the environmental persistence of *Leptospira* and highlight their potential role as ecological reservoirs in soil ecosystems.

## 2. Materials and Methods

### *Leptospira* strains and culture

*Leptospira interrogans* serovar Manilae strain L495 and *Leptospira biflexa* serovar Patoc strain Patoc I were cultured in Ellinghausen, McCullough, Johnson, and Harris (EMJH) liquid medium at 30 °C without agitation. Strains were subcultured twice weekly, grown to exponential phase and counted using a Petroff-Hausser cell-counting chamber (Hausser Scientific Company, Horsham, PA, USA) under dark-field microscopy. For co-incubation experiments, cultures were diluted in EMJH and concentration adjusted to reach a multiplicity of infection (MOI) of 50.

### Free-Living Amoebae strains and culture conditions

*Acanthamoeba castellanii* (Douglas) Page (ATCC 30010), *Dictyostelium discoideum* AX-3 (ATCC 28368) and *Hartmannella vermiformis* Page (ATCC 50236) were maintained under standard culture conditions in their respective growth media. *A. castellanii* was cultured in Peptone–Yeast–Glucose medium (PYG: Proteose peptone 20 g·L⁻¹, glucose 18 g·L⁻¹, yeast extract 2 g·L⁻¹, sodium citrate dihydrate 1 g·L⁻¹, MgSO₄·7H₂O 0.98 g·L⁻¹, Na₂HPO₄·7H₂O 0.355 g·L⁻¹, KH₂PO₄ 0.34 g·L⁻¹, Fe(NH₄)₂(SO₄)₂·6H₂O 0.02 g·L⁻¹; in water; pH 6.5). *D. discoideum* was grown in HL5 medium (Proteose peptone 5 g·L⁻¹, Thiotone E peptone 5 g·L⁻¹, glucose 10 g·L⁻¹, yeast extract 5 g·L⁻¹, Na₂HPO₄·7H₂O 0.35 g·L⁻¹, KH₂PO₄ 0.35 g·L⁻¹; in water; pH 6.5). *H. vermiformis* was cultured in medium 1033 (Bacto-peptone 10 g·L⁻¹, yeast extract 10 g·L⁻¹, yeast nucleic acid [RNA, type VI from Torula yeast] 1 g·L⁻¹, folic acid 15 mg·L⁻¹, hemin 1 mg·L⁻¹, phosphate buffer [KH₂PO₄ 18.1 g·L⁻¹, Na₂HPO₄ 25.0 g·L⁻¹] 20 mL·L⁻¹, heat-inactivated fetal bovine serum 10% v/v; in water; pH 6.5). All chemicals were of analytical grade and purchased from standard commercial suppliers.

All FLA cultures were maintained in T25 tissue culture flasks (Easy Flask, Nunc; Dutscher) at room temperature and subcultured two to three times per week by 1:25 dilution in a final volume of 5 mL. Cultures were used for experiments at 70–80% confluence.

Prior to infection assays, cells were harvested by centrifugation at 2,000*g* for 5 min at room temperature.

### Fluorescent labelling of *Leptospira/amoeba*

Exponentially growing *L. interrogans* or *L. biflexa* cultures were harvested by centrifugation (3,000*g*, 15 min, room temperature), washed in sterile phosphate-buffered saline, and resuspended at ∼10^8^ cells/mL in 1 mL of PBS containing 10 µM Vybrant™ CFDA-SE Cell Tracer (Ex/Em=494/521 nm, Thermo Fisher Scientific), a fixable fluorescein-based dye. Suspensions were incubated overnight at 30 °C in the dark and subsequently washed three times with PBS to remove excess dye. Where indicated, fluorescently labelled leptospires were fixed in 4% (w/v) paraformaldehyde (PFA) in PBS for 20 min at 30°C. Fixed bacteria were washed three times with PBS to remove residual PFA and prevent cytotoxic effects on FLA cells.

For confocal microscopy, amoebae were labelled prior to infection using CellTracker™ Red CMTPX (10 µM, 1H, Ex/Em = 586/614 nm; Thermo Fisher Scientific), a cell-permeable fluorescent dye or with Dapi (1µg/ml, 1H, Ex/Em = 358/461 nm; Thermo Fisher Scientific). Cells were then washed to remove excess dye and subsequently used for co-incubation experiments with fluorescently labelled leptospires.

### Co-incubation kinetic experiment

Amoebae were seeded in 12-well tissue culture plates (Costar®, clear TC-treated) at a density of 1 × 10⁵ cells per well and allowed to adhere for 1 h at room temperature in Peptone-Yeast (PY, PYG without glucose) medium. Where indicated, amoebae were pretreated with cytochalasin D (Sigma-Aldrich) at 20 µM for 1h to disrupt actin polymerization and inhibit actin-dependent uptake mechanisms.

Amoebae were then co-incubated with either live or PFA-fixed fluorescent *Leptospira* at a multiplicity of infection (MOI) of 50 (5 × 10^6^ bacteria per well), in a final volume of 1 mL, and incubated for 3 h at room temperature in the dark.

Following co-incubation, supernatants were removed and wells were gently rinsed three times with PBS. To eliminate extracellular bacteria, 1 mL of gentamicin solution (50 µg/mL final concentration) was added to each well and incubated for 2 h. After antibiotic treatment, gentamicin was removed and cells were washed three times with PBS.

Amoeba–*Leptospira* interactions were then assessed immediately after antibiotic treatment (t_0_) and at 1 h, 24 h, and 48 h post-infection using confocal microscopy, flow cytometry, and colony-forming unit (CFU) assays, as described in the corresponding sections.

### Confocal laser-scanning microscopy

For endpoint imaging of internalized leptospires, amoebae–*Leptospira* co-cultures were conducted in 12-well plates. A sterile round glass coverslip (No. 1.5 coverslip) was placed at the bottom of each well. Following co-incubation, samples were fixed in 4% PFA in PBS for 20 min at 30°C, washed, and mounted on glass slides with antifade mounting medium (ProLong™ Gold Antifade Mountant, Invitrogen). Stained samples were imaged on an inverted SP8 confocal laser-scanning microscope (Leica Microsystems) equipped with a white light laser tuneable between 470 and 670 nm and controlled by Leica Application Suite X (LAS X) software. Fluorescent leptospires (CFDA SE) were excited at 488 nm and emission was collected between ∼500–550 nm using hybrid detectors. Red CMTPX–stained amoebae were excited at 580 nm, and emitted fluorescence was collected between 610 and 620 nm. Images were acquired using a HC PL APO CS2 63×/1.40 NA or a HC PL APO CS2 40×/1.30 NA oil-immersion objective (Leica Microsystems). z-stacks were collected with a step size of 0.3–0.5 µm over a total depth of 6–10 µm, encompassing the entire trophozoites thickness.

Three-dimensional reconstructions and orthogonal views were generated directly in LAS X (3D module). Acquisition settings (laser power, detector gain, and offset) were kept constant within each experiment to allow qualitative comparison between conditions.

### Spinning-disk confocal live-cell imaging

Dynamic interactions between *Leptospira* and free-living amoebae were monitored by live-cell spinning-disk confocal microscopy. Amoebae (1 × 10⁵ cells) were seeded in 35 mm u-Dish (Ibidi) and allowed to adhere for at least 1 h in PY medium. CFDA-SE–labelled leptospires were then added (MOI=50), and dishes were immediately transferred to a Nikon Ti2-E inverted microscope (Nikon Instruments) equipped with a Yokogawa CSU-W1 spinning-disk unit, a motorized z-drive, and a sCMOS camera. The system was controlled by NIS-Elements software.

Fluorescent leptospires were excited with a 488 nm laser line and emission was collected through an appropriate GFP/FITC filter set. Unstained amoebae were visualized in parallel using transmitted-light widefield imaging to monitor cell morphology and motility. Images were acquired with a Plan APO 40×/0.95 NA or Plan Apo 60×/1.42NA immersion objective. At each time point, a z-stack spanning 20 µm in the z-axis was acquired with 1 µm spacing between optical sections. Time-lapse sequences were recorded over 2 to 5 min to capture initial contact, membrane interaction, and internalization events. Z-stacks were subsequently projected (maximum-intensity projection) and, where required, reconstructed in 3D to visualize the spatial relationship between leptospires and amoeba cytoplasm. Autofluorescence level background were assessed and signal to noise ratio was evaluated over 10:1 (Fig S1).

### Image processing and quantification

All images were processed using LAS X (Leica) or NIS-Elements (Nikon) softwares and, when needed, further edited in FIJI/ImageJ. Only linear adjustments of brightness and contrast were applied equally across the entire image and within each experimental set. Intracellular vs. extracellular leptospires were assessed by inspection of individual z-slices and orthogonal reconstructions. For quantitative analyses (e.g. percentage of infected amoebae), at least 30 fields containing 20 cells were analyzed in each condition for 5 independent experiments using identical acquisition parameters.

### Flow cytometry analysis

CFDA-SE-labeled leptospires were incubated with amoebae. Following incubation, adherent amoebae were detached by gentle flushing and pipetting, collected, and directly resuspended in 1 mL of 4% paraformaldehyde (PFA). Cells were then fixed at 30°C for 20 min. Fixed samples were centrifuged at 2,000*g* for 5 min resuspended in 500 µL of 1X PBS and transferred to flow cytometry tubes. Samples were stored at 4 °C and protected from light prior to acquisition on the BD FACSCanto™ II Clinical Flow Cytometry System. After doublet exclusion, amoebae were gated based on forward and side scatter (FSC/SSC), and CFDA-SE fluorescence was acquired in the FITC channel (Fig S2). Positive fluorescence threshold was set using amoeba in the absence of leptospires. Following instrument calibration, up to 20,000 events were recorded per condition. Data were analyzed using FlowJo™ v10 software.

The percentage of CFDA-SE-positive amoebae was calculated as: 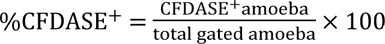.

As a control for phagocytic activity, amoebae were incubated with 1.0 µm red fluorescent FluoSpheres™ polystyrene beads (580/605; Thermo Fisher), detected using the 488 nm laser and acquired in the PE channel (Fig S3).

### Determination of bacteria viability

Viable *Leptospira* were quantified by colony-forming unit (CFU) enumeration on semisolid EMJH agar. To quantify extracellular bacteria, 200 µL of culture supernatant from each well were collected and serially diluted up to 10⁻⁵ (two successive 1:100 dilutions followed by one 1:10 dilution). Aliquots of 100 µL from each dilution were plated onto EMJH agar and incubated at 30 °C. Plates were examined every two days until visible colonies appeared. Plates showing no growth after 6 weeks of incubation were considered negative. To assess the viability of intracellular *Leptospira*, amoebae were harvested from the same wells, washed three times with PBS to remove residual extracellular bacteria, and mechanically lysed using sterile glass beads in a MagnaLyser instrument (Roche Diagnostics; two cycles of 90 s at 2,000 rpm). A 200 µL aliquot of the resulting lysate was serially diluted and plated onto EMJH agar under the same conditions as supernatant samples.

### Isolation of *Leptospira* from environmental soil samples

Authorization to collect environmental samples was obtained from the South Province of New Caledonia (Arrêté n°1689-2017/ARR/DENV). To investigate the potential co-occurrence of members of the *Leptospira* genus and FLA, six moist soil and one water samples were collected at the Parc zoologique et Forestier in Nouméa in May 2020. For each site, 5 g of topsoil were collected using a soil core sampler (3 cm diameter, 5–7 cm depth). Samples were collected by direct coring into sterile 15 mL polypropylene tubes and processed within 2 h of collection. Each tube was vigorously shaken with 5 – 10 mL of sterile water to resuspend the soil. After sedimentation of large particles for 5–15 min, 2 mL of the supernatant were either (i) filtered through a sterile 0.45 µm membrane filter into a tube containing 2.5 mL of 2× EMJH medium or (ii) kept unfiltered when longer sedimentation times were allowed. To limit bacterial and fungal contamination, 500 µL of 10× concentrated STAFF selective supplement (sulfamethoxazole 400 µg/mL, trimethoprim 200 µg/mL, amphotericin B 50 µg/mL, fosfomycin 4 mg/mL and 5-fluorouracil 1 mg/mL) (25) were added to each culture. Tubes were incubated at 30 °C and monitored daily by dark-field microscopy for the presence of spirochetes. When contaminating microorganisms were observed, cultures were filtration through a 0.45 µm membrane filter and re-supplemented with STAFF and fresh EMJH. Upon detection of spirochetes, cultures were serially diluted and plated on semisolid EMJH agar. Volumes of 50 µL and 200 µL were spread at different dilutions and incubated at 30 °C. Subsurface colonies characteristic of *Leptospira* appeared in positive samples within 3–10 days. One to five morphologically typical isolated colonies per plate were selected, presence of leptospires was confirmed by dark-field microscopy and clonally subcultured in liquid EMJH medium to obtain pure clonal isolates.

### Identification of *Leptospira* isolates by MALDI-TOF mass spectrometry

*Leptospira* cultures in exponential growth phase were harvested by centrifugation (1 mL culture), washed once in PBS, and subjected to a standard water–ethanol protein extraction. Pellets were subsequently processed by formic acid/acetonitrile extraction by resuspension in 15 µL of 70% formic acid and 15 µL of acetonitrile, followed by centrifugation at 11,000 x g for 3 min. After centrifugation, 1 µL of the resulting supernatant was spotted onto a MALDI-TOF target plate, air-dried, and overlaid with α-cyano-4-hydroxycinnamic acid (HCCA) matrix.

For each isolate, four independent spots were prepared and each spot was read twice. Spectra were acquired in linear positive mode over a mass range of 0–10,000 m/z (laser frequency 60 Hz). Raw spectra were normalized and baseline-corrected before comparison with an in-house *Leptospira* reference database currently covering 69 *Leptospira* species. Species or clade assignments were determined based on spectral similarity scores.

### Isolation of free-living amoebae from environmental soil samples

Soils samples were transferred to sterile 15 mL polypropylene tubes within 2 h of collection and vigorously shaken with 5–10 mL of sterile distilled water to resuspend soil particles. After brief sedimentation of coarse material, the soil suspensions were sequentially filtered through 20 µm and 6 µm pore-size membrane filters to enrich for free-living amoebae (FLA).

The 6 µm pore-size filters were then placed upside down onto Petri dishes containing Page Amoebic Saline (PAS) agar (NaCl 0.12 g·L⁻¹, MgSO₄·7H₂O 0.004 g·L⁻¹, CaCl₂·2H₂O 0.004 g·L⁻¹, Na₂HPO₄ 0.142 g·L⁻¹, KH₂PO₄ 0.136 g·L⁻¹, agar 1.5% w/v), previously seeded with 200 µL of a suspension of *Enterobacter aerogenes* in PAS medium inactivated by heat treatment (95 °C, 20 min) to serve as a bacterial food source.

Plates were incubated at room temperature and monitored regularly by light microscopy for the emergence and migration of trophozoites. Upon visible proliferation of FLA, agar fragments containing amoebae were excised and transferred onto fresh PAS agar plates to obtain progressively enriched cultures. Repeated subculturing steps were performed until stable amoeba growth was achieved. The successive stages of amoeba isolation and subculture are shown in Fig S4.

### Putative identification of amoebae using 18S rRNA gene and ITS sequencing

Genomic DNA was extracted from amoeba cultures originating from environmental samples using the QIAamp DNA Mini Kit (QIAGEN), according to the manufacturer’s instructions, and eluted in 30 µL of AE buffer. DNA concentration and purity were assessed by spectrophotometry (NanoDrop 2000, Thermo Fisher Scientific).

To determine the taxonomic identity of environmental amoebae, partial regions of the 18S rRNA gene and the internal transcribed spacer (ITS) were amplified by PCR using amoeba-specific primers. Amplifications were performed on a LightCycler system using the FastStart SYBR Green I Master Mix (Roche). Primer pairs Ami6F1 (5′-CCAGCTCCAATAGCGTATATT-3′) and Ami9R (5′-GTTGAGTCGAATTAAGCCGC-3′) were used to amplify a fragment of the 18S rRNA gene, while primers JITSFW (5′-GTCTTCGTAGGTGAACCTGC-3′) and JITSRV (5′-CCGCTTACTGATATGCTTAA-3′) were used for amplification of the ITS region.

PCR products were verified by electrophoresis on 1.2% agarose gels, excised when necessary, and purified using the MinElute PCR Purification Kit (QIAGEN). Purified amplicons were sequenced using the ABI Prism BigDye Terminator v3.1 Cycle Sequencing Kit (Applied Biosystems). Sequencing reactions were carried out on a GeneAmp 9700 thermocycler and analyzed by capillary electrophoresis on an ABI 3130xl Genetic Analyzer (Applied Biosystems).

Resulting sequences were assembled, quality-checked, and compared against reference sequences in the NCBI nucleotide database using BLAST.

### Statistical analysis

Statistical analyses were performed using GraphPad Prism software (version 10.0.0, GraphPad Software, San Diego, CA, USA). Data are presented as mean ± SEM unless otherwise stated. Prior to statistical testing, datasets were assessed for normality using the Shapiro–Wilk test. For comparisons between paired experimental conditions (before vs after treatment), parametric tests were applied when data followed a normal distribution (paired two-tailed Student’s t-test). When normality was not met, non-parametric Wilcoxon signed-rank tests were used. A p-value < 0.05 was considered statistically significant.

The number of biological replicates and independent experiments for each analysis is indicated in the corresponding figure legends.

## 3. Results

### *Leptospira* are internalized within environmental *Acanthamoeba castellanii*

To determine whether *Leptospira* can be internalized by free-living amoebae, fluorescently labeled bacteria (Fig 1A) were incubated with *A. castellanii* trophozoites and monitored by confocal microscopy. In the absence of bacteria, only amoebae cytoplasm was visible and no fluorescent signal was detected (Fig 1B). In contrast, incubation with *L. interrogans* resulted in a rapid association of bacteria with amoebae surface and subsequent internalization, visualized as discrete green, fluorescent foci within the cytoplasm (Fig 1C). By 1 h of incubation, a large proportion of amoebae displayed intracellular fluorescent leptospires (Fig 1D). Similar internalization patterns were observed with the saprophytic, non-pathogenic species *L. biflexa*, which also localized within trophozoites (Fig 1E and Fig 1F).

**Fig 1.**
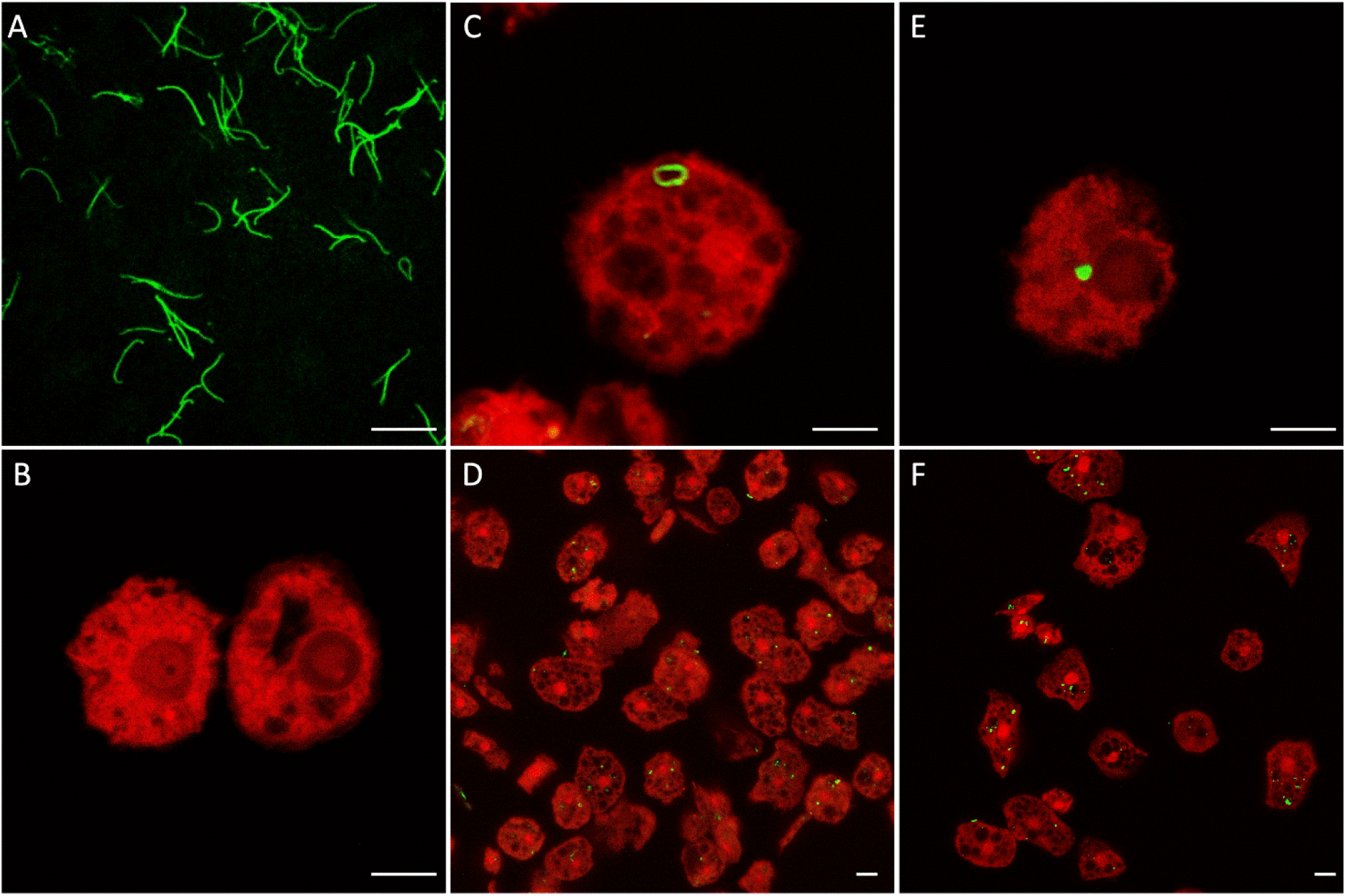
Internalization of *Leptospira* by *Acanthamoeba castellanii*. Confocal micrographs of (A) Fluorescently labeled *L. interrogans* prior to contact with amoebae, showing typical helical morphology. (B) Negative control: *A. castellanii* trophozoites incubated without bacteria, stained in red. (C) Early stage of internalization: a single *L. interrogans* (green) associated with the amoeba cytoplasm (red). (D) Multiple trophozoites after 5 h of co-incubation, showing numerous internalized bacteria. (E–F) Internalization of *L. biflexa* into *A. castellanii* trophozoites, observed as green intracellular foci within amoebal cytoplasm. Scale bar: 10 µm.

Three-dimensional optical sectioning and reconstruction (Movie S1) confirmed the intracellular localization of leptospires. Live-cell confocal imaging further revealed the dynamics of early interactions (Movie S2, S3 and Fig S5), capturing bacteria in close contact with amoebae plasma membrane before internalization. Notably, this process did not involve classical phagocytic structures, as no phagocytic cups were observed. Entry was accompanied by a rapid morphological shift of leptospires from their characteristic filamentous shape to a rounded, possibly vacuolated form once inside the host cell.

### Classical phagocytic mechanisms partially contribute to *Leptospira* internalization by amoeba

Because the entry of *Leptospira* into *A. castellanii* did not display the hallmarks of canonical phagocytosis, we examined whether internalization was mediated by actin-dependent endocytic pathways. Amoebae were pretreated with Cytochalasin D, a potent inhibitor of actin polymerization that disrupts cytoskeleton-dependent uptake. As expected, Cytochalasin D treatment impaired amoebae motility and induced a round morphology (Fig 2A), consistent with cytoskeletal disruption. Under these conditions, the proportion of infected amoebae was reduced by approximately twofold (from 68.6% to 30.6% for *L. interrogans*, and from 74% to 25.5% for *L. biflexa,* Fig 2B). Although significant, this reduction did not abolish internalization, indicating that translocation of leptospires into amoebae is only partially dependent on actin-driven endocytic processes and may also involve alternative entry pathways.

**Fig 2.**
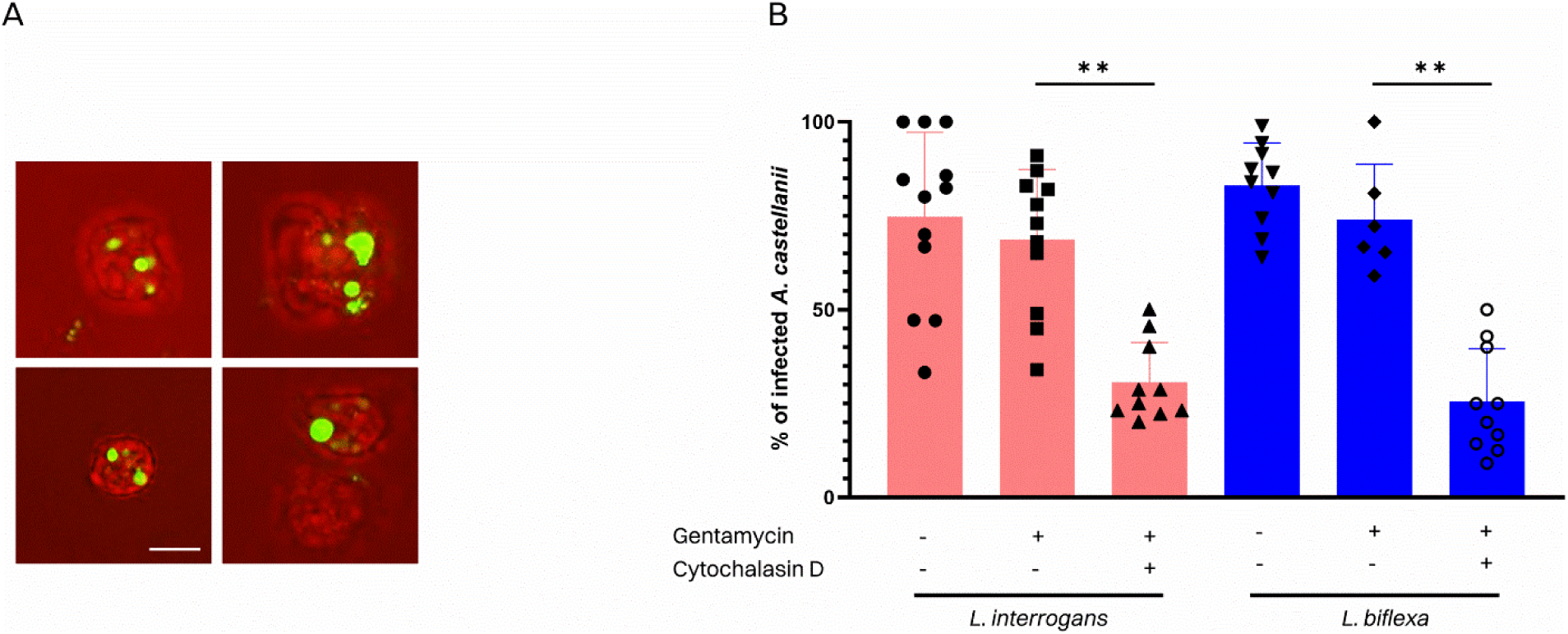
Partial dependence of *Leptospira* internalization on actin cytoskeleton dynamics. (A) Representative confocal images of *A. castellanii* trophozoites incubated with fluorescently labeled *Leptospira* (green) after Cytochalasin D treatment. Amoeba are shown in red. Internalized leptospires appear as discrete green foci within the amoeba cytoplasm. (B) Quantification of the proportion of infected *A. castellanii* following co-incubation with *L. interrogans* (pink) or *L. biflexa* (blue). Pretreatment of amoebae with Cytochalasin D significantly reduced bacterial uptake for both species. Bars represent mean ± SEM of independent experiments; individual data points are shown. Scale bar: 10 µm. (**p < 0.01).

### Viable leptospires actively contribute to their internalization

To investigate whether *Leptospira* entry into amoebae involves an active, bacteria-driven process, we performed co-incubation assays using live or PFA-fixed *L. interrogans* and *L. biflexa* labelled with CFDA-SE. This approach allowed to discriminate between active internalization and passive uptake by amoebae. Prior to infection, we verified that PFA treatment did not alter leptospiral fluorescence intensity: live and fixed bacteria displayed comparable CFDA-SE signals by flow cytometry (Fig S6). In parallel, flow cytometric analysis (FSC/SSC) revealed no changes in amoeba’s size or granularity after incubation with PFA-treated leptospires, indicating preserved morphology.

Internalization was quantified by flow cytometry following gentamicin protection (Fig 3). We performed this analysis with three free-living amoebae: *A. castellanii*, *D. discoideum*, and *H. vermiformis* as recognized hosts for a broad range of human bacterial pathogens. All three species were capable of internalizing *Leptospira*, although with markedly different efficiencies (Fig 3). *H. vermiformis* consistently exhibited the lowest internalization rates for both *L. interrogans* and *L. biflexa* (26.1% and 12.5% respectively). In contrast, *A. castellanii* and *D. discoideum* internalized *L. interrogans* at substantially higher levels (above 68%). In contrast, saprophytic *L. biflexa* showed substantially lower internalization levels than pathogenic *L. interrogans* across both amoeba hosts (Fig 3A and Fig 3B).

**Fig 3.**
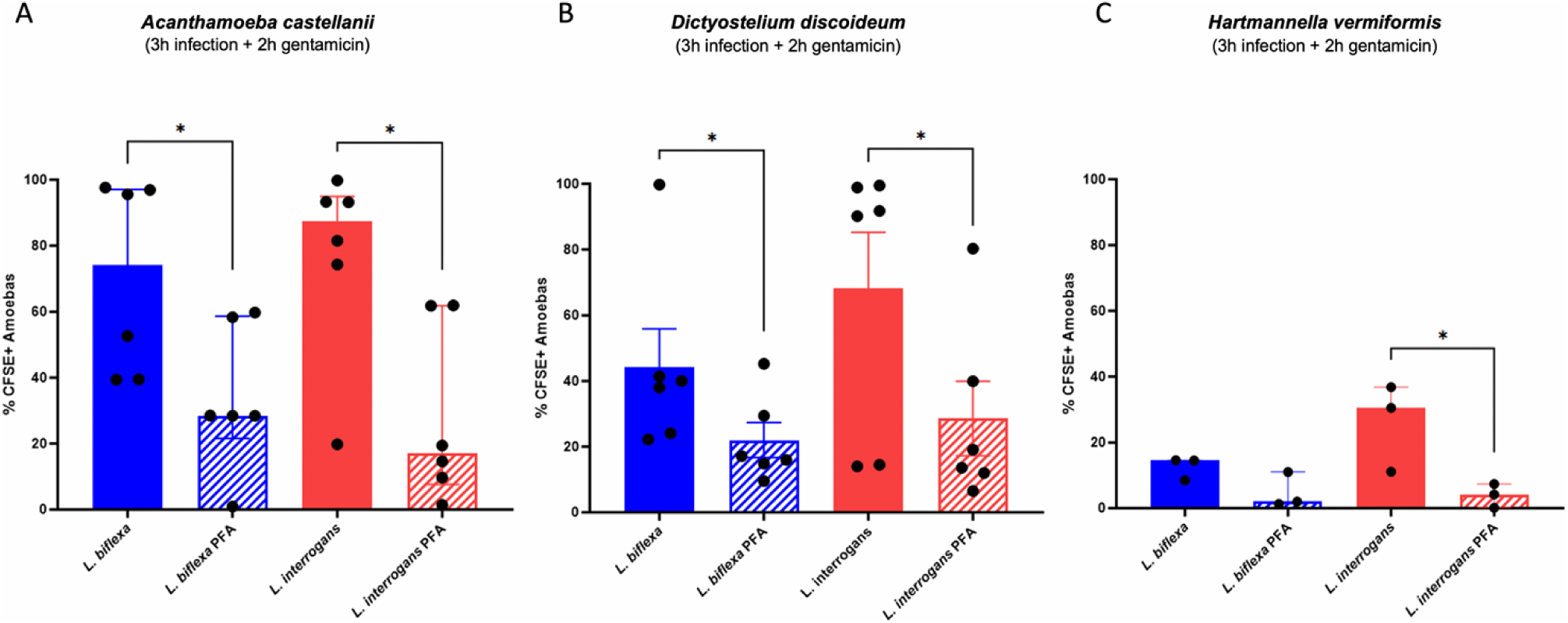
Internalization of *Leptospira* by environmental amoebae requires bacterial viability. Flow cytometry analysis of amoebae infected with live or paraformaldehyde (PFA)-killed *Leptospira* (3 h infection + 2 h gentamicin treatment). Infection rates were determined as the percentage of CFDE-SE positive amoebae. (A) *Acanthamoeba castellanii*, (B) *Dictyostelium discoideum*, and (C) *Hartmannella vermiformis*. Live bacteria are represented by solid bars, whereas PFA-fixed bacteria are represented by hatched bars. In all three species, uptake was reduced for PFA-killed bacteria compared to live bacteria, for both *L. interrogans* (red) and *L. biflexa* (blue). Bars represent median with interquartile range (IQR) of independent experiments. (*p < 0.05; **p < 0.01; ***p < 0.001).

Pre-fixation of leptospires with 4% PFA resulted in a significant reduction in the proportion of CFDA-SE-positive amoebae in most host-bacteria combinations, indicating that bacterial viability contributes to efficient internalization. However, this effect did not reach statistical significance for *L. biflexa* in *H. vermiformis* (Fig 3). Although internalization was not completely abolished by bacterial fixation, the consistent decrease in internalization observed with non-viable bacteria supports a model in which leptospires physiological activity contributes to entry into amoebae.

### *Leptospira* survive within amoebae but do not proliferate

We next examined whether *Leptospira* can persist intracellularly in FLA, potentially exploiting them as “Trojan horses” for survival. To address this, infection dynamics were monitored in *A. castellanii* using both confocal microscopy, flow cytometry and CFU, combined with gentamicin protection assays to eliminate extracellular bacteria (Fig4).

At the early stage of infection, approximately 70% of *A. castellanii* cells contained fluorescently labeled leptospires, with no significant difference between *L. interrogans* and *L. biflexa* (Fig 4A and Fig 4B). By 24 h of incubation, the proportion of infected amoebae had significantly decreased to 16.5% and 24.1% for *L. interrogans* and *L. biflexa*, respectively (Fig 4B). Similar values were observed at 24 and 48 h. These results indicate that both species of *Leptospira* can persist intracellularly in *A. castellanii* for at least 48 h. However, the absence of increase in the infection rates suggests an absence of bacterial replication within amoebae. Confocal imaging also showed no evidence of newly released, free-swimming bacteria that would indicate a bacterial exit from amoeba.

**Fig 4.**
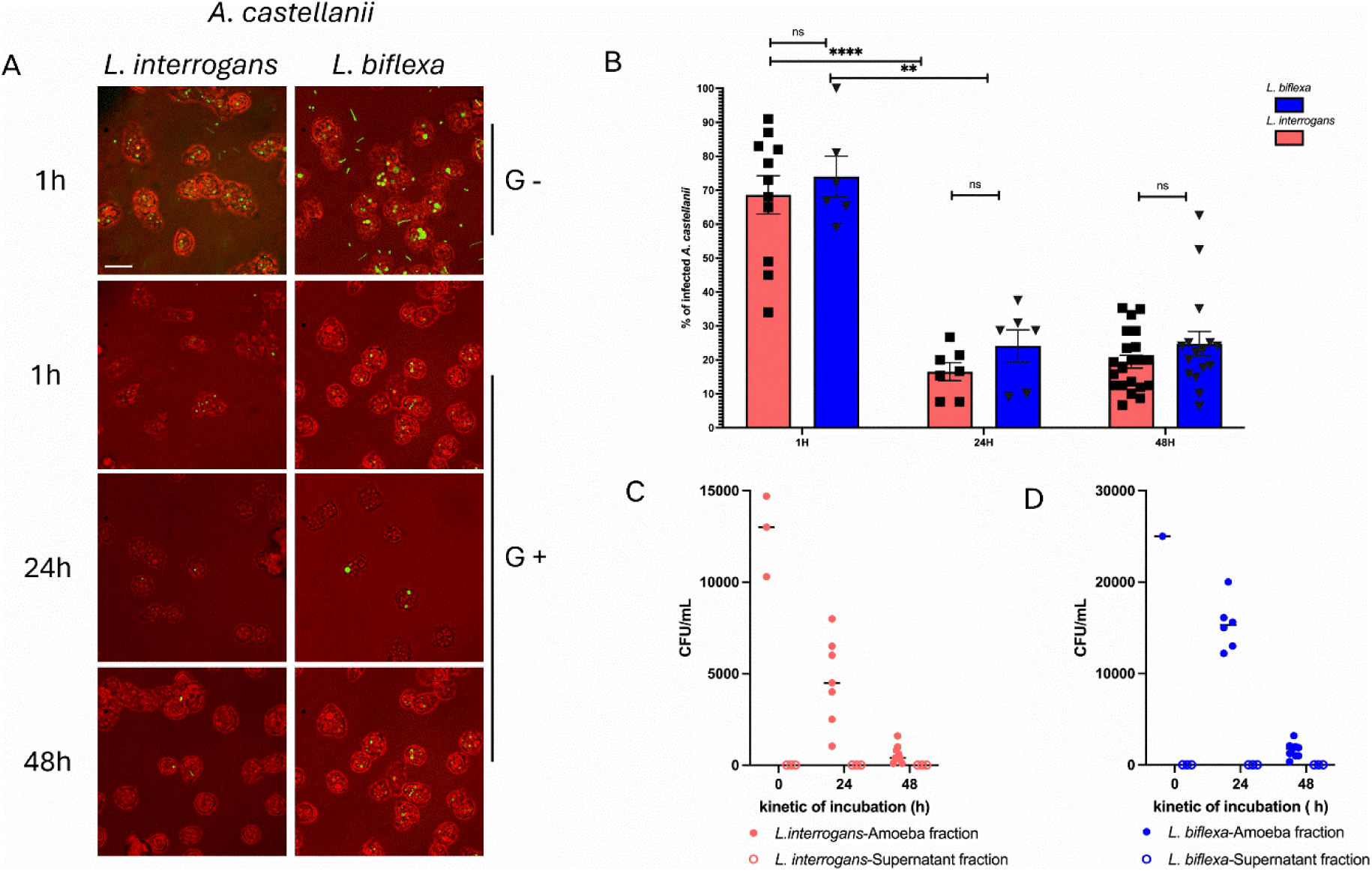
Interaction of *Leptospira interrogans* and *Leptospira biflexa* with *Acanthamoeba castellanii*. (A) Representative fluorescence microscopy images of *A. castellanii* incubated with *L. interrogans* or *L. biflexa* for 1 h, 24 h, and 48 h. Amoebae were stained with CellTracker™ Red CMTPX (red), and leptospires are shown in green. Panels labeled G− correspond to non-gentamicin-treated conditions, whereas G+ indicates samples treated with gentamicin to eliminate extracellular bacteria prior to imaging. Scale bar: 20 µm (B) Quantification of the percentage of infected amoebae at 1 h, 24 h, and 48 h post-incubation after gentamicin treatment. Bars indicate mean ± SEM. (ns: not significant; **p < 0.01; ****p < 0.0001). (C–D) Viable bacterial counts (CFU/mL) recovered from the amoeba fraction and corresponding supernatant at 0 h, 24 h, and 48 h for (C) *L. interrogans* and (D) *L. biflexa*. Intracellular-associated bacteria progressively declined over time without evidence of intracellular replication, while supernatant fractions showed minimal recoverable CFUs after gentamicin treatment.

Because the detection of fluorescent signal within amoebae does not directly prove bacterial viability and could reflect progressive digestion, we assessed colony-forming ability as a proxy for survival. At 0, 24, and 48 h post-infection, both culture supernatants and amoebae lysates were plated on semi-solid EMJH medium. No colonies were recovered from culture supernatants at any time point, confirming that gentamicin treatment effectively eliminated extracellular leptospires and indicating that internalized bacteria did not subsequently escape from amoebae during the incubation period (Fig 4C and 4D). In contrast, colony-forming units were consistently recovered from amoebae lysates up to 48 h post-infection (Fig 4C and 4D), indicating that a fraction of internalized leptospires remained viable and culturable despite the marked change in cell morphology changes. Importantly, CFU counts progressively declined over time, supporting the conclusion that although leptospires persist within amoebae for at least 48 h, they do not proliferate intracellularly.

### *Leptospira* interact with FLA environmental soil isolates

To assess whether the interactions observed in vitro with model amoebae could be relevant in natural environments, we analyzed seven soils samples collected in New Caledonia, a French island territory in the Pacific region. *Leptospira* spp. were detected in all seven soil samples. Pure *Leptospira* isolates were successfully obtained from six samples. Free-living amoebae isolation was attempted in six samples and was successful in all cases, confirming the co-occurrence of *Leptospira spp*. and free-living amoebae within the same habitats. MALDI-ToF identification of *Leptospira* isolates revealed representatives of the saprophyte S1 and pathogenic P2 clades, while 18S rRNA (V4–V5 and ITS) sequencing of amoebae isolates putatively identified *Naegleria spp*, and *Acanthamoeba spp* (Table I).

**Table I.** Co-isolation and molecular identification of *Leptospira* spp. and FLA from environmental soil samples.

| Sample Name | Leptospira identification |  | FLA identification |  |  |  |  |
| --- | --- | --- | --- | --- | --- | --- | --- |
|  | Presence of *Leptospira. spp* (Liquid culture) | *Leptospira* identification (Maldi ToF score ; clade) | Isolation of FLA | Fragment | Length (pb) | Blast results | Match % |
| Soil-P1-00 | + | n/d | + | V4-V5<br>ITS | 632<br>479 | *Negleria gruberi*<br>*Acantamoeba clarki* | 100%<br>95.59% |
| Soil-PZF20-01 | + | unmatched *L. sp*<br>1.90-2.00 ; S1 clade | + | V4-V5<br>ITS | 673<br>753 | *Naegleria. Sp*<br>*no match* | 100%<br>/ |
| Soil-PZF20-02 | + | *L. levettii*<br>2.42-2.57 ; S1 clade | + | V4-V5<br>ITS | 763<br>774 | *Acantamoeba lenticulata*<br>*Acantamoeba castellanii* | 98.95%<br>96.61% |
| Soil-PZF20-03 | + | unmatched *L. sp*<br>1.79-1.91 ; S1 clade | + | V4-V5<br>ITS | 633<br>487 | *Negleria clarki*<br>*Negleria clarki* | 100%<br>100% |
| Soil-PZF20-04 | + | unmatched *L. sp*<br>1.87-2.00 ; P2 clade | n/d | n/a | n/a | n/a | n/a |
| Soil-PZF20-05 | + | *L. jelokensis*<br>2.26-2.49; S1 clade | + | V4-V5<br>ITS | 516<br>495 | *Acantamoeba lenticulata*<br>*Naegleria. Clarki* | 100%<br>95.59% |
| Water-PZF20-06 | + | *L. levettii*<br>2.42-2.57 ; S1 clade | + | V4-V5 | 778 | *Acantamoeba lenticulata* | 100% |

To investigate whether environmental free-living amoebae could internalize *Leptospira*, co-incubation assays were performed using these soil-derived FLA and fluorescently labeled *L. interrogans*. Fluorescence microscopy showed intracellular localization of leptospires across all tested wild amoebae isolates (Fig 5). At early stages (1 h), bacteria were observed within amoebae cytoplasm, and by 24 h intracellular fluorescent leptospires were still detected, confirming sustained persistence within non-axenic, soil-derived FLA, thereby providing a model that more closely reflects natural environmental conditions.

**Fig 5.**
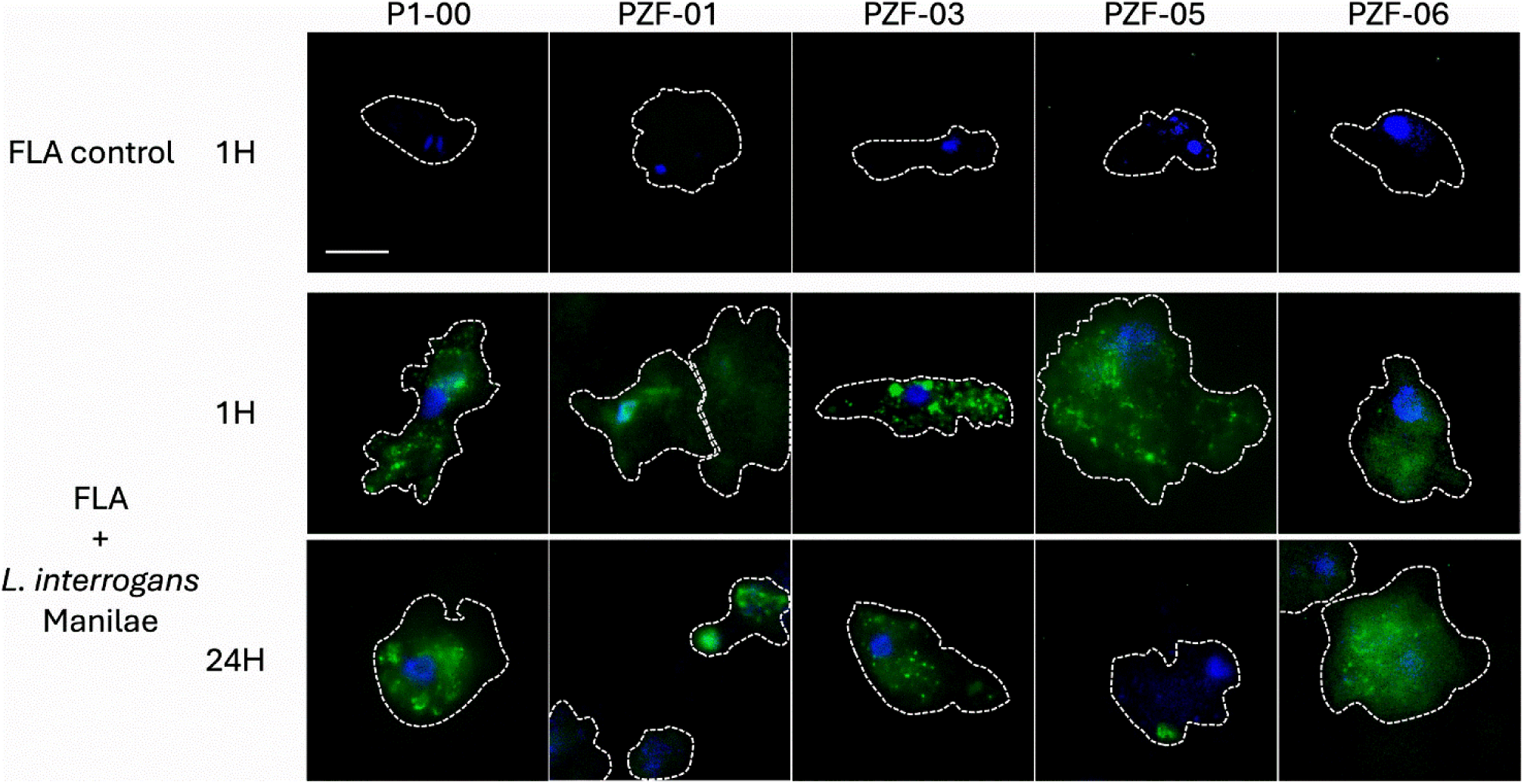
Intracellular localization of fluorescent *Leptospira interrogans Manilae* within FLA environmental soil isolates. Representative confocal fluorescence microscopy images showing intracellular localization of CFDA-SE–labeled *Leptospira* (green) within amoebae. Amoeba nuclei were stained with DAPI (blue), and cell contours are delineated by dashed white lines to highlight cell boundaries. Upper panels show uninfected control amoebae showing absence of green fluorescence. Middle and lower panels reveal amoebae incubated with fluorescent *Leptospira interrogans Manilae*, displaying discrete intracellular green fluorescent signals distributed throughout the cytoplasm. Images were acquired as optical sections by confocal laser-scanning microscopy. Scale bar: 10 µm

These observations extend our previous findings obtained with laboratory strains and demonstrate that naturally occurring environmental free-living amoebae are also permissive to *L. interrogans* Manilae internalization and intracellular persistence.

## 4. Discussion

Free-living amoebae are recognized as environmental reservoirs for bacteria, providing protective and nutrient-rich microenvironments that can support survival and, in many cases, intracellular replication. *Legionella pneumophila*, *Mycobacterium tuberculosis*, *Salmonella enterica*, and *Helicobacter pylori* exemplify pathogens that exploit amoebae as “Trojan horses”, persisting within their cytoplasm and often exhibiting increased resistance to disinfectants and environmental stresses (24). In this study, we investigated whether *Leptospira*, a pathogen well known for its ability to persist in soils and waters (2,4,26), engages in comparable interactions with FLA, and whether such interactions could contribute to its environmental life cycle.

Using confocal microscopy and live-cell imaging, we demonstrate that both pathogenic (*L. interrogans*) and saprophytic (*L. biflexa*) leptospires are rapidly internalized by *Acanthamoeba castellanii*. Internalization occurred within the first hour of co-incubation and resulted in discrete intracellular bacteria, as confirmed by three-dimensional reconstruction. Notably, entry did not display canonical hallmarks of phagocytosis, such as the formation of phagocytic cups, and was accompanied by a pronounced morphological transition of leptospires from their characteristic helical shape to a rounded intracellular form. Together, these observations suggest that *Leptospira* entry into amoebae does not rely exclusively on classical phagocytic pathways and may involve non-canonical uptake mechanisms. Pharmacological disruption of amoeba actin cytoskeleton further supports this interpretation: Cytochalasin D treatment significantly reduced, but did not abolish, *Leptospira* internalization, indicating that actin-driven phagocytosis contributes to uptake but is not strictly required. This partial dependence contrasts with the predominantly phagocytic uptake described for many amoeba-associated bacteria (18,27) and points to a more complex interaction in which *Leptospira* may exploit additional, actin-independent processes.

Consistent with this hypothesis, we show that bacterial viability influences internalization efficiency. Live leptospires were internalized by three phylogenetically and ecologically distinct FLA (*A. castellanii*, *D. discoideum*, and *H. vermiformis*) at significantly higher rates than PFA-fixed bacteria, except for *H. vermiformis* with *L. biflexa*. Moreover, *A. castellanii* and *D. discoideum* exhibited higher overall internalization levels compared to *H. vermiformis*. However, no statistically significant difference was observed between pathogenic *L. interrogans* and saprophytic *L. biflexa* within each host. These findings indicate that internalization is largely amoeba-dependent under our experimental conditions, although leptospires viability also contributes to the process. PFA fixation cross-links bacterial surface proteins, potentially masking microbe-associated molecular patterns and impairing amoeba recognition independently of viability. In addition, fixation abolishes motility, a recognized leptospiral virulence-associated trait that may enhance contact with amoebae; therefore, reduced internalization cannot be attributed solely to a loss of leptospires metabolic activity. Finally, fixation increases bacterial rigidity and reduces membrane flexibility, which may hinder engulfment by amoeba pseudopodia. The lower internalization of fixed leptospires likely reflects a combination of impaired recognition, loss of motility, and altered physical properties, rather than a single mechanism.

Elucidating the molecular determinants of entry and intracellular fate will therefore require additional approaches, such as comparative uptake assays using motility-impaired leptospires, targeted inhibition of candidate surface structures, or transcriptomic and proteomic analyses of bacteria during early amoeba contact. These strategies would help disentangle the relative contributions of bacterial motility, surface architecture, and host recognition pathways.

Once internalized, *Leptospira* were able to persist within amoebae for up to 48 hours without evidence of intracellular replication. Although the proportion of infected FLA cells declined substantially over time, viable bacteria were consistently recovered from amoeba lysates at late time points. In contrast, no *Leptospira* (CFUs) were recovered from culture supernatants at any stage, indicating an absence of detectable escape of culturable leptospires from FLA under our experimental conditions, although the presence of bacteria in a viable but non-culturable state cannot be excluded. Collectively, these findings support a model in which amoebae permit transient intracellular survival of *Leptospira* without acting as amplification hosts.

Similar intracellular persistence without replication has been reported for other environmental bacteria (28) and is often interpreted as a state of dormancy or reduced metabolic activity that favors survival under adverse conditions (29). Whether *Leptospira* can withstand amoeba encystment remains an important open question, as encystment could further enhance environmental persistence as shown for other bacterial pathogens (18).

The ability of *Leptospira* to persist intracellularly for up to 48 hours suggests that amoebae may serve as short-term shelters during the environmental cycle. This time window could allow leptospires to withstand transient stressors such as nutrient limitation, desiccation, ultraviolet radiation, or exposure to antimicrobial compounds. Future studies should determine whether amoebae actively confer protection to leptospires under such harsh environmental conditions, and whether leptospires can persist within amoebae during encystment, which could further enhance their environmental resilience. Given the vast diversity of soil protists, it is conceivable that other protist hosts may support long-term intracellular persistence, or even allow limited multiplication under specific ecological conditions, although this remains to be demonstrated. Nevertheless, we did not observe bacterial release into the supernatant, letting leptospires exit mechanism unresolved. Possible scenarios include release following amoeba death, gradual leakage below detection thresholds, or transmission upon predation or physical disruption of amoebae in natural environments. These hypotheses warrant targeted investigation.

Strikingly, these observations parallel the interaction of *Leptospira* with mammalian macrophages (20). Both *L. interrogans* and *L. biflexa* can survive ingestion by human and murine macrophages without apparent intracellular replication, with bacterial loads progressively declining over time. Amoebae and macrophages are both professional phagocytes that rely on conserved cellular modules for chemotaxis, phagocytosis, oxidative stress responses, and metal-dependent antimicrobial mechanisms. Survival strategies effective in one context may translate to the other. Given that free-living amoebae pre-date macrophages evolutionarily, it is plausible that persistence mechanisms observed in mammalian immune cells originated from ancient environmental interactions, shaping *Leptospira*’s ability to transiently evade host defenses. Importantly, saprophytic Leptospira species are considered evolutionarily more ancient than pathogenic lineages (30), suggesting that environmental lifestyles preceded vertebrate host adaptation. In this context, the ability to transiently survive within phagocytic cells may have initially evolved in saprophytic strains as a response to predation by environmental protozoa. The observation that *L. biflexa* exhibits intracellular survival dynamics broadly comparable to *L. interrogans* supports the idea that this phenotype represents an ancestral ecological adaptation trait rather than a virulence determinant acquired later in the pathogenic clades. It is therefore conceivable that mechanisms allowing tolerance to intracellular stress were “trained” in environmental saprophytes, subsequently retained and possibly refined in pathogenic lineages, where they may contribute to partial evasion of macrophage-mediated clearance. Finally, the co-isolation of FLA and *Leptospira* from New Caledonian soils reinforces the ecological relevance of our findings. The ability of soil-derived amoebae to internalize *L. interrogans* under laboratory conditions indicates that these interactions are not restricted to reference strains or artificial systems. Even low-level persistence of viable pathogenic *Leptospira* in soils can sustain transmission, and transient protection within amoebae may be sufficient to bridge periods of unfavorable environmental conditions. This framework helps understanding frequent environmental detection of *Leptospira* despite their apparent fragility under laboratory conditions and highlights free-living amoebae as overlooked, yet potentially important, components of leptospirosis ecology.

Even in the absence of intracellular multiplication, transient survival within FLAs may therefore provide a selective advantage by extending environmental viability and increasing the likelihood of host encounter, contributing to the persistence of *Leptospira* in natural ecosystems. Beyond this short-term shelter, it remains possible that interactions with other free-living amoebae or protists could result in extended intracellular persistence or even support limited multiplication under specific ecological conditions.

In conclusion, this study demonstrates that free-living amoebae, including *Acanthamoeba castellanii*, *Dictyostelium discoideum*, *Hartmannella vermiformis* and natural isolates, can internalize both pathogenic and saprophytic *Leptospira* through a process partially dependent on actin-mediated phagocytosis and requiring bacterial viability. Once internalized, leptospires persist for at least 48 hours without replicating, maintaining viability but showing progressive decline in intracellular load. These observations reveal that amoebae can transiently harbor *Leptospira* providing a temporary intercellular refuge that may contribute to survival in harsh natural soil and water environments.

The co-occurrence of *Leptospira* and free-living amoebae in natural soils and the capacity of environmental isolates to internalize the pathogen strengthen the ecological relevance of this interaction. By bridging environmental microbiology and host-pathogen interactions, this study provides new insight into the ecological cycle of *Leptospira*. It identifies free-living amoebae as previously underappreciated components of the environmental stage of leptospirosis transmission.

## Supporting Information Legends

**Movie S1. Three-dimensional reconstruction of intracellular *Leptospira* within FLA.**

3D reconstruction generated from confocal z-stack images showing fluorescently labeled *Leptospira* localized within the cytoplasm of FLA, confirming intracellular bacterial localization.

**Movie S2. Live imaging of Leptospira internalization by FLA.**

Live spinning-disk confocal microscopy showing close contact between a fluorescent *Leptospira* and a FLA followed by bacterial internalization into the host cell.

**Movie S3. Dynamic internalization and morphological change of *Leptospira* inside FLA.**

Live Spinning-disk confocal microscopy showing the interaction of a fluorescent *Leptospira* with an FLA, its subsequent internalization, and the rapid morphological transition of the bacterium following entry into the host cell.

## Author contributions

Conceptualization: CG and RT. Writing-original draft preparation: AL and RT. Investigation: AL, AA, CI, EM and RT. Methodology: AL, AA, CI and RT. Data analysis: AL, AA, CI and RT. Validation: AL, AA, CI, EM, VBS, MP, CG and RT. All authors contributed to the review and editing of the manuscript and approved the submitted version.

## Acknowledgments

We gratefully acknowledge our scientific colleagues and collaborators for their valuable discussions and contributions to the experimental work. We thank Marie-Estelle Soupé-Gilbert and Malia Kainiu for maintaining amoeba and *Leptospira* strains, as well as for performing amoeba 18S rRNA gene sequencing and MALDI-TOF identification of *Leptospira* strains. We also thank Alexandre Giraud-Gatineau for valuable scientific discussions and insights. We are grateful to Viviane Chenal-Francisque for providing the amoeba strains.

---

**Fig S1.**
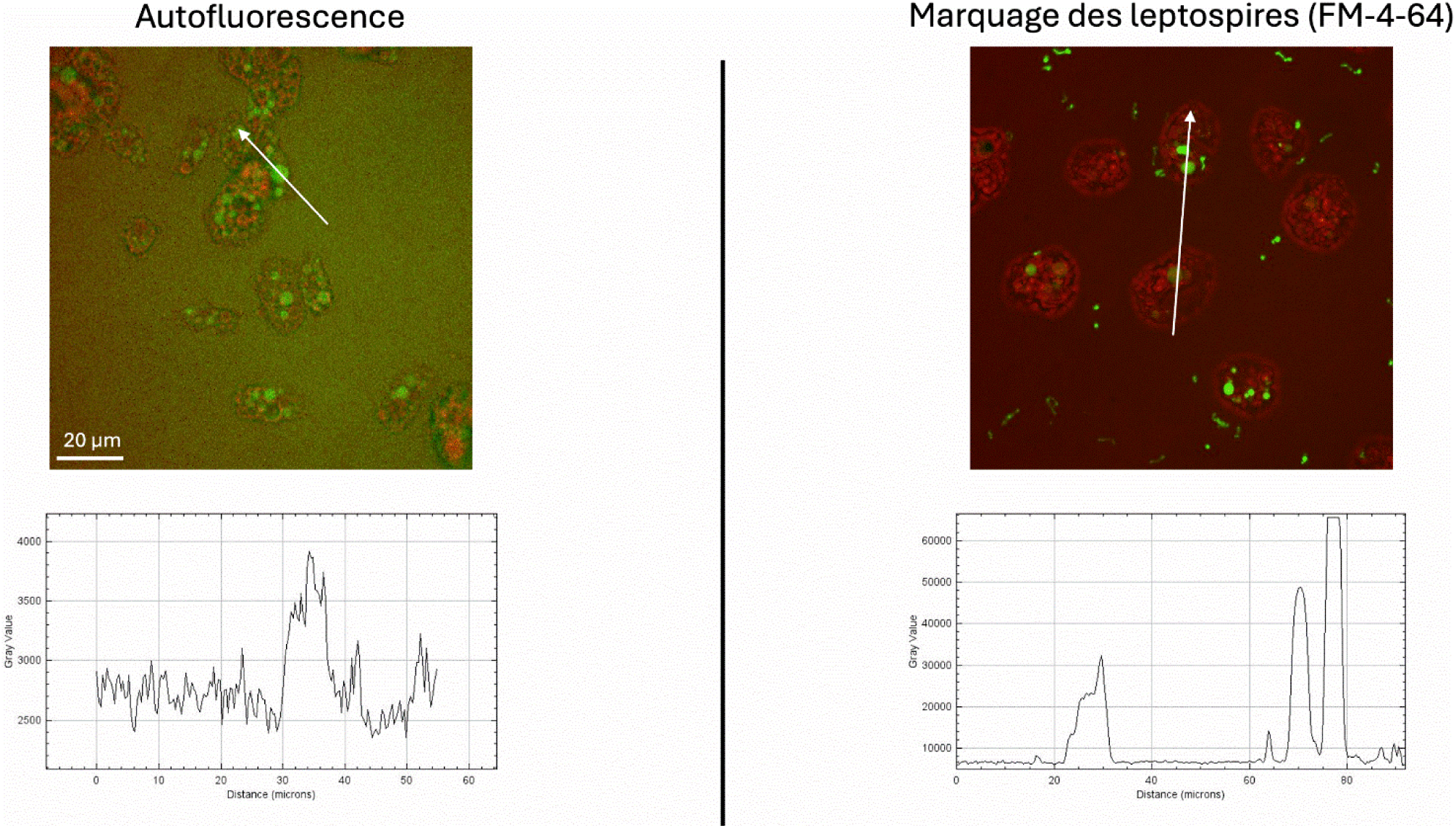
Live spinning-disk confocal imaging and discrimination of *leptospira* signal from amoeba autofluorescence. Live-cell spinning-disk confocal microscopy was performed to distinguish true leptospiral fluorescence from background autofluorescence. Left panel (Autofluorescence control): Amoebae imaged in the absence of specific bacterial membrane staining. Green signal corresponds to intrinsic cellular autofluorescence. The white arrow indicates the region used for line-scan intensity analysis. The corresponding fluorescence intensity profile (bottom graph) shows low intensity (Max 4000 A.U.), low-amplitude, diffuse background signal across the scanned distance. Scale bar: 20 µm. Right panel (CFDA-SE –labeled leptospires): Amoebae co-incubated with leptospires stained with the membrane dye FM4-64. Discrete green fluorescent structures corresponding to bacteria are visible within amoebae. Line-scan analysis (bottom graph) reveals sharp, high-intensity peaks (Max 66000 A.U.), clearly distinguishable from autofluorescence background levels observed in control conditions

**Fig S2.**
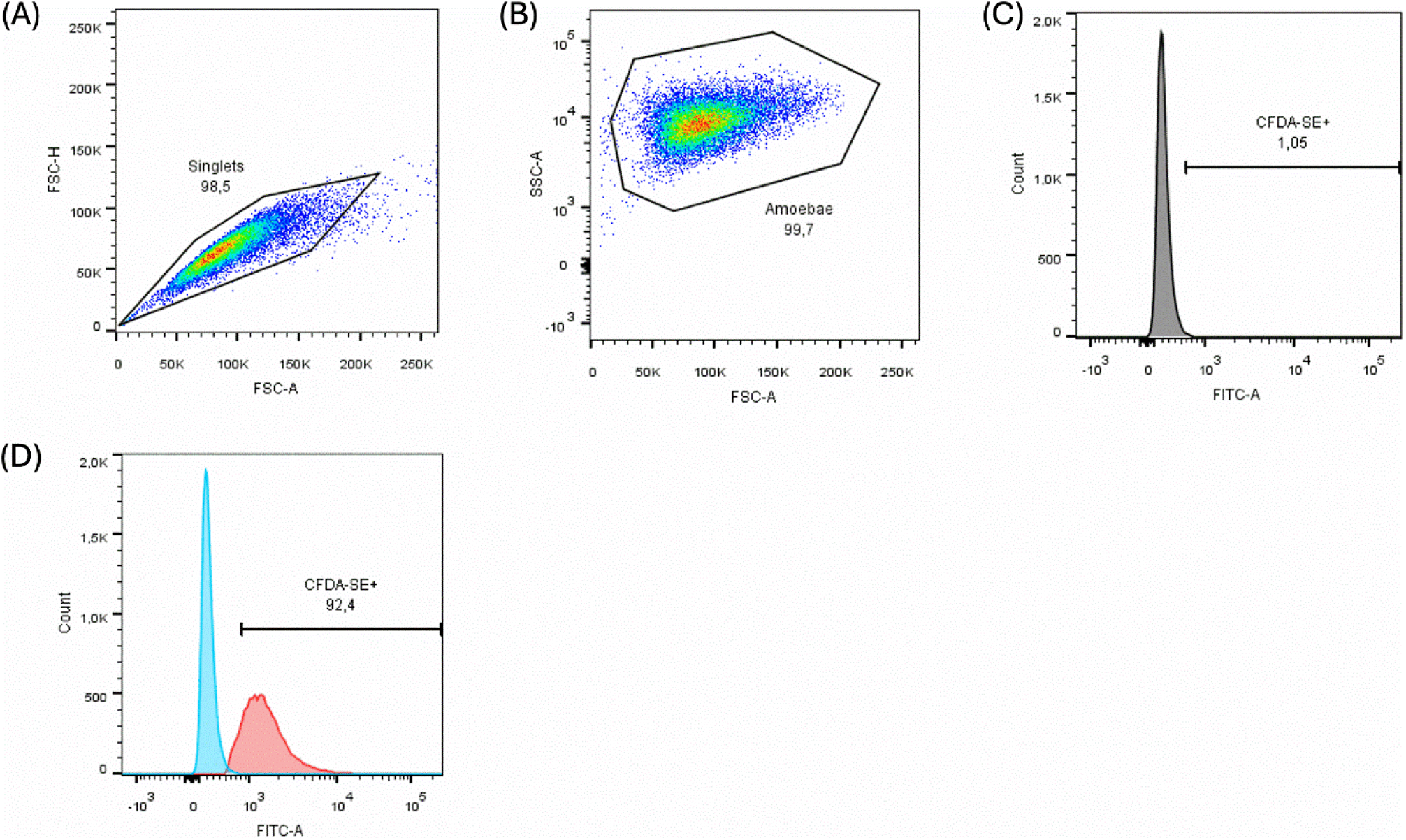
Flow cytometry gating strategy used to quantify CFDA-SE-positive amoebae. (A) Doublets were excluded using forward scatter area versus height (FSC-A/FSC-H) parameters, and single-cell events (singlets) were retained for further analysis. (B) Amoebae were identified and gated according to their forward and side scatter properties (FSC-A/SSC-A), allowing exclusion of debris and non-cellular events. (C) The CFDA-SE positivity threshold was defined on FITC-A histograms using non-infected control samples, with the gate set to include approximatively 1% of events. (D) Representative overlay of FITC-A histograms from non-infected (blue) and infected (red) amoebae showing the application of the same CFDA-SE+ gate. The percentage of CFDA-SE-positive amoebae was calculated relative to the total gated amoeba population.

**Fig S3.**
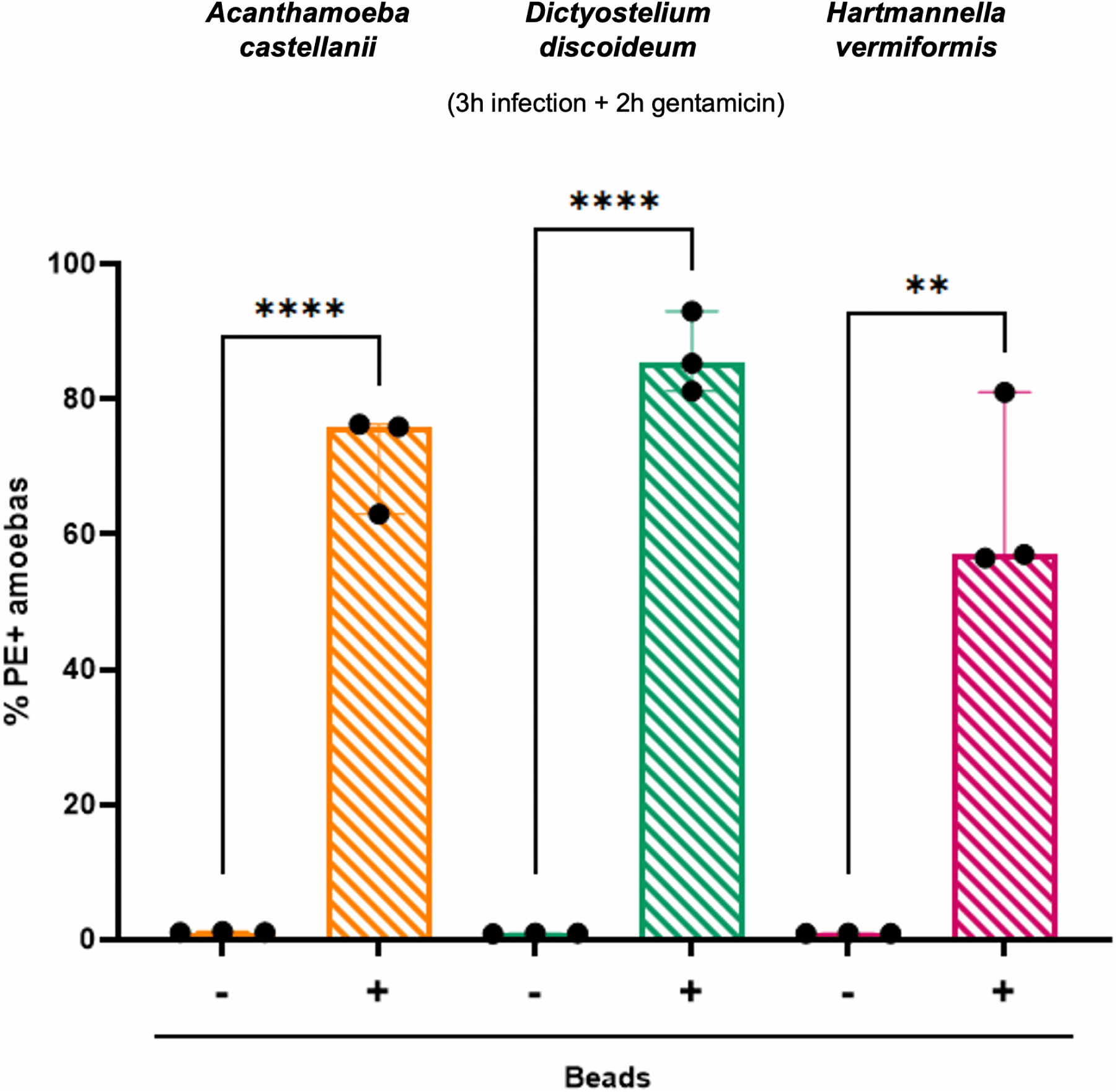
Amoebae’s phagocytosis capacity. Percentage of PE-positive amoebae following internalization of fluorescent beads after 5 h of co-incubation (3 h exposure followed by 2 h gentamicin treatment) at an MOI of 50. Bead internalization was assessed in *Acanthamoeba castellanii*, *Dictyostelium discoideum* and *Hartmannella vermiformis*. Bars represent mean values and dots indicate individual biological replicates. Statistical significance between control (- beads) and bead-exposed (+ beads) conditions is indicated.

**Fig S4.**
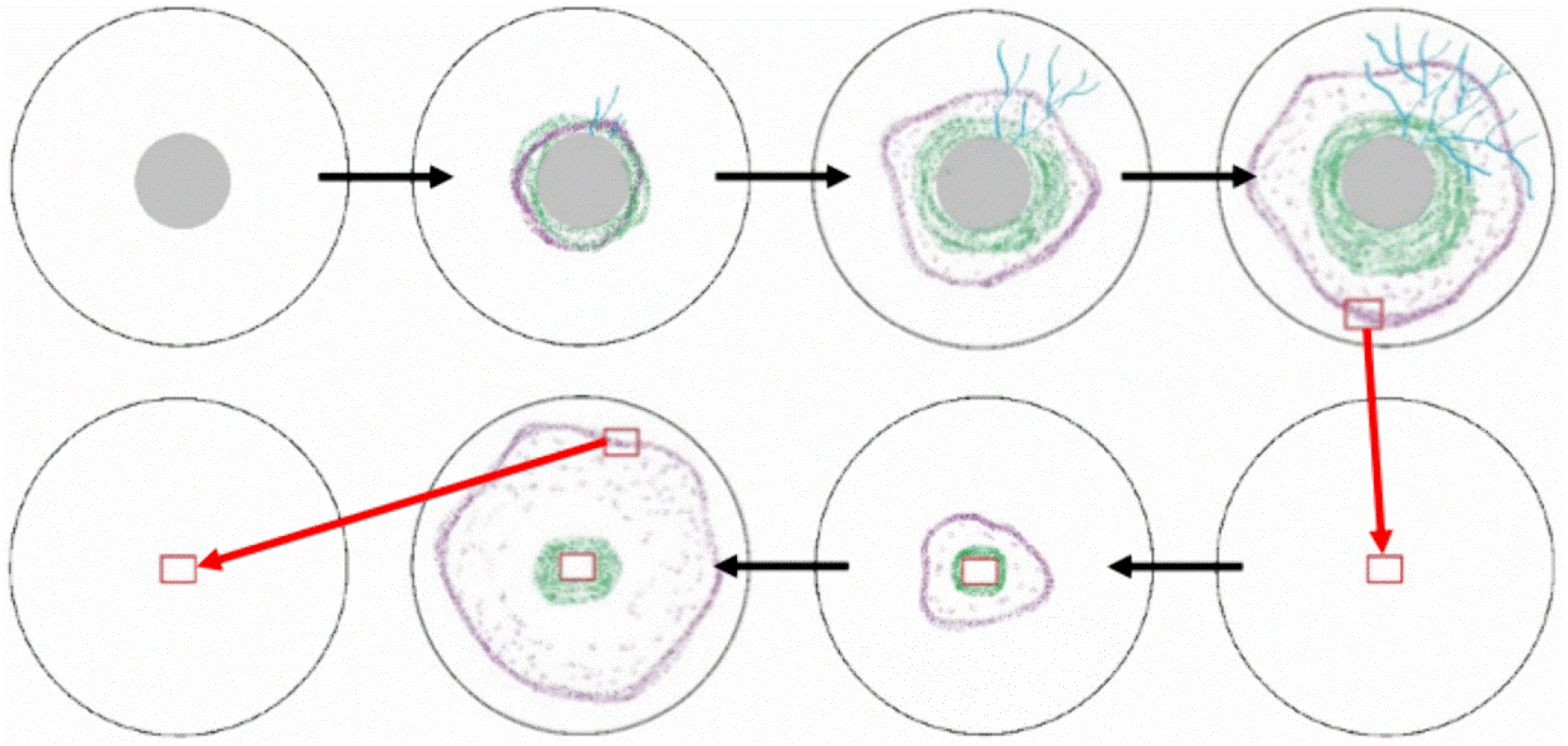
Workflow for isolation of free-living amoebae from environmental samples. Schematic representation of the successive steps used to isolate FLA on non-nutrient agar plates. Environmental samples were filtered (grey membrane) and the filters were placed onto agar plates seeded with heat-inactivated bacteria. Over time, amoebae (purple) migrate from the filter and proliferate, while contaminant bacteria (green) and fungal elements (blue) may also be present. When amoeba growth was observed, a defined piece of agar containing trophozoites (red rectangle) was excised and transferred to a fresh plate to obtain purified cultures.

**Figure S5.**
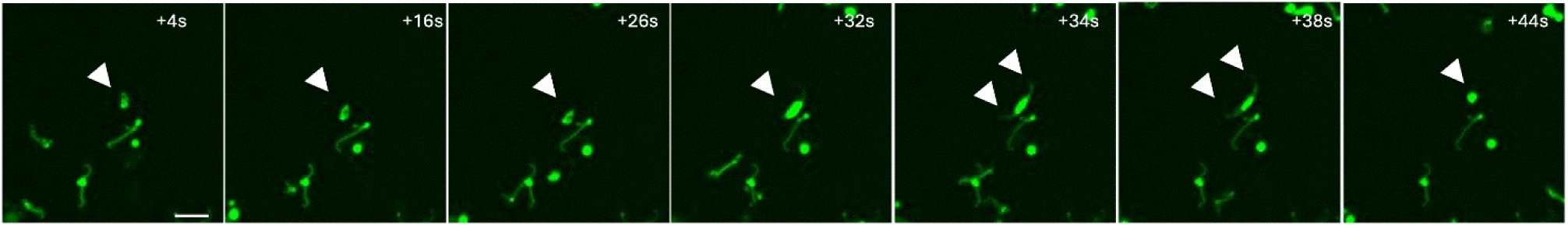
Time-lapse spinning-disk confocal imaging of live fluorescent *Leptospira*. Representative sequential frames from live-cell spinning-disk confocal microscopy showing CFDA-SE–labeled *Leptospira* over time (indicated in seconds). White arrowheads highlight individual bacteria tracked across successive frames. The series illustrates active motility and dynamic morphological changes, including transitions from elongated helical forms to more compact or rounded conformations after internalization. Scale bar: 5 µm.

**Fig S6.**
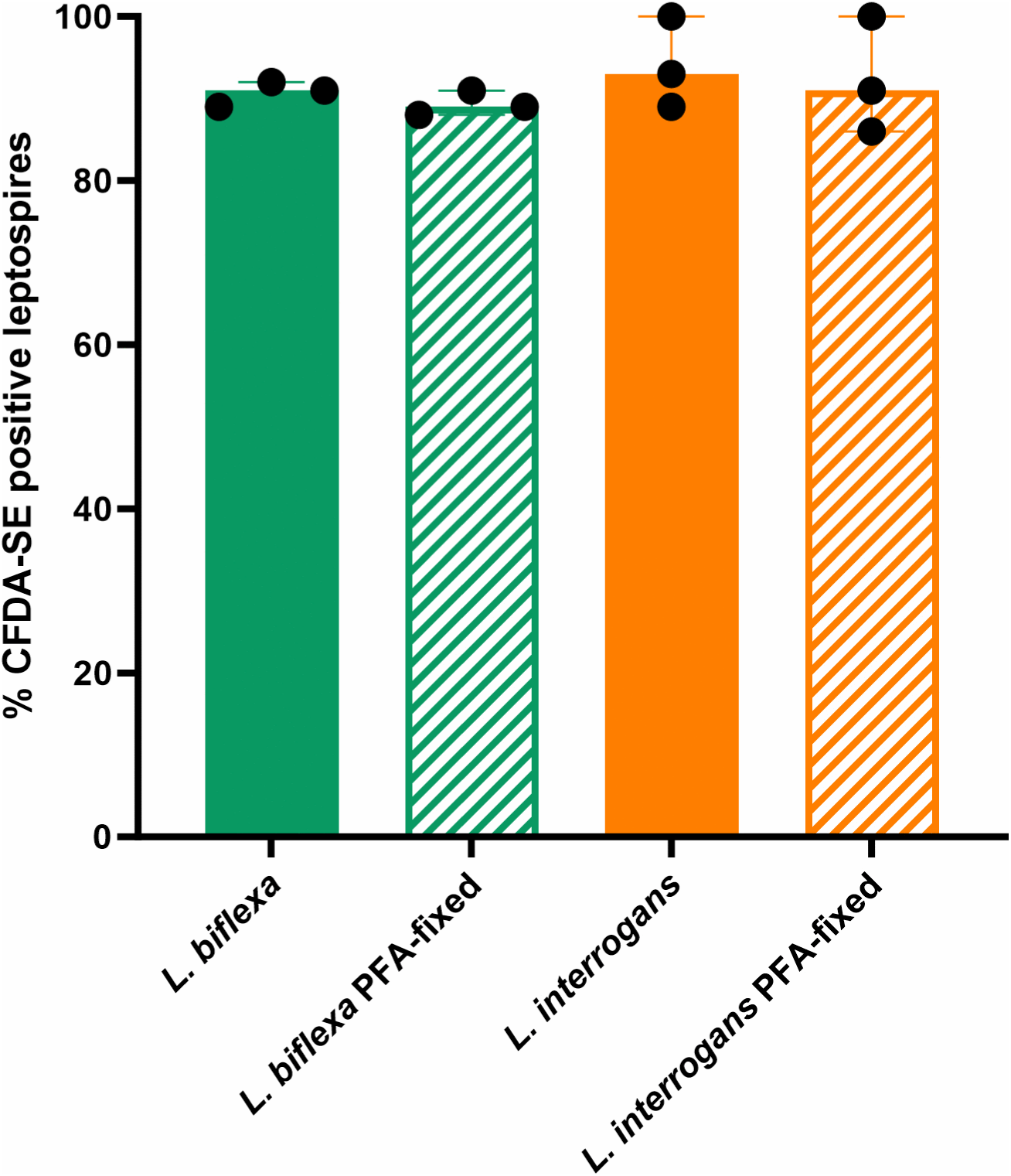
Percentage of CFDA-SE–labeled viable and PFA-fixed leptospires. Percentage of CFDA-SE–positive *Leptospira biflexa* and *Leptospira interrogans*, either viable or fixed with 4% paraformaldehyde (PFA), as determined by flow cytometry. Comparable fluorescence levels were observed between live and PFA-fixed bacteria for both species, indicating that paraformaldehyde treatment did not alter CFDA-SE fluorescence intensity.

